# Dual-lipid metabolomics revealed the communication of MTB or MB with Bovine alveolar macrophages in lipid metabolisms

**DOI:** 10.1101/2021.07.06.451399

**Authors:** Weifeng Gao, Yurong Cai, Gang Zhang, Xiaoping Wang, Juan Wang, Yong Li, Yujiong Wang

## Abstract

*M. tuberculosis(*MTB) and *M. bovis*(MB) of the Mycobacterium tuberculosis complex (MTBC) are the causative agents of the notorious infectious disease tuberculosis(TB) in a range of mammals, including cattle and human. The lipid composition of MTB/MB performed imperative function as invading host macrophage. In the present study, a dual-lipid metabolomics were used to elucidate the differences in lipid composition of MTB and MB and the different responses in lipid metabolisms of **bovine alveolar macrophage** challenged by MTB/MB. The lipid metabolomics of MTB and MB indicated that there were significant differences in lipid composition of both bacteria that the level of various lipids belonged to Glycerophospholipids, Sterol Lipids, Fatty Acyls and Polyketides exhibited differences between MTB and MB. Meanwhile, both MTB and MB with different lipid composition could invoked different responses in lipid metabolisms of the host macrophage. MTB infection mainly induced the increase in content of Polyketides and Glycerophospholipids in macrophages, whereas MB infection induced the level of Glycerophospholipids and Sterol Lipids of macrophages. Furthermore, we identified TAG 13:0-18:5-18:5 of MTB and PC(16:1(9E)/0:0), PI(20:2(11Z,14Z)/22:6(4Z,7Z,10Z,13Z,16Z,19Z)), 4,6-Decadiyn-1-ol isovalerate and LacCer(d18:1/24:1(15Z)) of MB caused the different variations in lipid metabolisms of macrophage following MTB/MB attacks, respectively. Finally, we proposed MTB and MB with different lipid compositions could successfully colonize in macrophage by different mechanisms that MTB could promote the formation of foam cells of macrophage for its colonization and development, while MB mainly through suppressing the macrophage autophagy to escape the immune responses of host.

**Importance:** The differences in lipid composition of MTB and MB and the different responses in lipid metabolisms of **bovine alveolar macrophage** challenged by MTB/MB were elucidated in this study. The lipid composition of MTB and MB exhibited significantly different patterns, which could induced different responses of the host macrophage during the infection process of MTB or MB, respectively. MTB infection mainly induced the increase in content of Polyketides and Glycerophospholipids in macrophages, whereas MB infection induced the level of Glycerophospholipids and Sterol Lipids of macrophages.The results presented here thus provides a comprehensively dual-lipid metabolomics profile in MTB/MB and MTB/MB-attacked macrophages, which deepened our understanding of the interaction between host and pathogens in lipid metabolisms.

## INTRODUCTION

As global health problem, Tuberculosis affects millions of people annually and is the main cause of death in some developing countries and regions (WHO). Tuberculosis **(TB)** is characterized by tuberculous granulomas and calcified necrotic lesions in tissues and organs, which results in cough and even death in both patients and animals. Statistically, there were 10 million people worldwide being infected with TB in 2017, and 1.6 million died due to such notorious disease (1). Tuberculosis is a chronic infectious disease caused by the Mycobacterium tuberculosis complex (MTBC) which mainly comprises *M. tuberculosis*(**MTB**) and *M. bovis*(**MB**) and cause tuberculosis (TB) in a broad range of mammalian species, including humans and bovine (2–5). According to the statistics that about 10% of the 10 million cases of human TB are caused by MB infection, leading a tremendous economic burden through production loss and control costs (Heath et al., 2013)(5–8). The species in MTBC show greater than 99% nucleotide sequence similarity, indicating a clear host preference, and it has been proved by experimental bovine infection models that MTB seems unable to be maintained in non-human animal populations (i.e., through the cycle of infection, disease, and transmission).

As an imperative immune cells in vivo, Macrophages play a key role in host immunity against infection (9). Macrophages are not only the parasitic sites of MTBC, but also the most important immune cells against MTBC infection. In the process of MTBC infection, macrophages can regulate the host inflammatory response and immune response through phagocytosis, antigen presentation and secretion of various cytokines (10). Previously, series studies showed that macrophages activated Toll like receptor (TLR), transforming growth factor-β (TGF-β), Hippo (HPO) and other cellular signaling pathways of host macrophages after being infected by MTB, and further eliminated MTB through apoptosis or autophagy (11–13). Meanwhile, during MTBC infection, macrophages can provide nutrients and a place for survival and reproduction for MTBC, and lipids of macrophage functioned as the main carbon source for MTBC development(8, 11). Researchers have shown that MTBC infection can cause alterations in lipid metabolisms of macrophages, leading to significant increase in content of intracellular lipid and cholesterol and contributing the tranformation of macrophage to foam cells(14). And the cell foaminess of macrophages will weaken the bacteria-clearing function of macrophages in the host immune response, thus conducive to the infection and colonization of MTBC (15). Remarkably, the foaming of macrophages is closely related to the lipid substances specific to MTBC. It has been proved that MTB can secrete lipid mycoacid to inhibit the autophagy reaction of mouse macrophages, and cause the imbalance of lipid metabolism leading to macrophage foaming**(16)**. Additionally, for successful colonization, MTBC could avoid proteolytic hydrolysis and subsequent immune responses by preventing phagosome acidification and the fusion of phagosomes and lysosomes(17, 18). It is worth noting that lipids, as the main component of the cell wall of mycobacterium, accounting for about 60% of the dry weight, are also indispensable in the process of MTB escaping from the immune system of host macrophages. However, the lipid composition of MTBCs were different. The virulence of MTBC mainly depends on the composition of large amount of lipids contained in the cell wall that the MTBC with the higher lipid content exhibits stronger virulence. Studies have shown that MTB could synthesis phthiocerol dimyco - ceroserate (PDIM) lipid on its cell wall to masking the pathogen associated molecular patterns of macrophages PAMPs(19). This mechanisms lead to a host cell toll-like receptors cannot properly recognize pathogenic MTBC and the downstream of the MyD88 and Nf-kappa B signal pathway responses cannot be activated, which resulted in no activation in immune responses of host macrophages (19). However, the distinct lipid composition of the main pathogenic microbial MTB and MB were still unclear, and whether MTB or MB with different lipid composition could induce different alterations in lipid metabolisms of macrophage were still unknown. Hence, elucidating the host responses of lipid metabolisms to MTB and MB will undoubtedly contribute our understanding of host-MTBC communication, which is helpful for reducing or even eliminating the risk of human TB infection.

In this study, the lipid composition of macrophage following MTB or MB and both pathogenic microbial MTB and MB were determined by an in-depth lipid metabolomics analysis. We found that the lipid composition of MTB and MB exhibited significantly different patterns, which could induced different responses of the host macrophage during the infection process of MTB or MB, respectively. Indeed, the infection of both MTB and MB with distinct lipid composition resulted in different alterations in lipid metabolisms of macrophages. Ultimately, through integrated network analysis on the communication between lipid composition of infectious macrophages together with MTB and MB identified the hub lipids of MTB and MB. These specific lipids functioned in the mechanisms of which caused the different variations in lipid metabolisms of macrophage in response to MTB or MB attacks. This present study explored the differences in lipid composition and macrophage responses to MTB and MB, and elucidated these differences in lipid composition lead to different mechanisms of early infection in the host.

## MATERIALS AND METHODS

### Bacterial strains culture

There are three virulent strains of M. bovis and three virulent strains of M. tuberculosis. Three M. bovis strains MB1,MB1,MB3 were collected and stored in Key lab of Ministry of Education for protection and utilization of special biological resources in western China.Three M. tuberculosis strains Mtb1,Mtb2,Mtb3 were obtained from The Fourth People’s Hospital of Ningxia Hui Autonomous Region.The six strains were grown in Middlebrook 7H9 medium supplemented with 10% oleic acid albumin dextrose catalase (OADC) enrichment (Difco BD, Franklin Lakes, NJ), 0.5% glycerol and 0.05% Tween 80. Before every infection experiment, Bacteria were harvested at mid-log phaseh point and resuspended in Dulbecco’s modified eagle medium (DMEM) medium supplemented with 10% fetal bovine serum (complete medium).

### Isolation and infection of bovine alveolar macrophages

This study was approved by the ethics committee for use and care of animals at Ningxia University (Yinchuan,China). Three unrelated, age-matched male calves were selected from a tuberculosis-free herd and total bovine alveolar macrophage were harvested by pulmonary lung lavage via tracheal infusion of physiological saline solution. The cell suspension was washed by centrifugation,then discard the supernatant and add 3 times the volume of the erythrocyte lysate to the lysate. Gently mix by pipetting, and leave on ice for 10 min. Centrifuge the supernatant at 800 rpm for 5 minutes, discard the upper red supernatant, collect the pellet, wash it with PBS, and centrifuge at 800 rpm for 5 minutes. Approximately 4 × 10^7^ total lung cells from each animal were cultured in AIM VTM Medium(research grade,12055091,Thermo Fisher) supplemented with 10% foetal bovine serum (FBS) and 4× antibiotics (Penicillin&Streptomycin&Amphotericin B solution), γ-Irradiated[Biological Industries] for 16 h at 38 °C,5% CO2 .Wash twice with 38°C pre-warmed PBS and change to AIM VTM Medium with 10%FBS no antibiotics in second day before infection. The purity of alveolar macrophages obtained by this method can reach 95%(5, 20). Bovine alveolar macrophage were infected with M.bovis strains or M.tuberculosis strains at a multiplicity of infection (MOI) of 10 bacilli per alveolar macrophage.Each group of cells of treatment groups were challenged with Bacteria (10:1) multiplicity of infection (MOI) for 4h and 6 h respectivelyat 38 °C, with 5% CO2.We performed uninfected control ,MB-infected cells and M.tb-infected cells at two time points (4, 6 h), totaling 18 samples.All MTB or MB infection experiments were conducted in the biosafety level 2 (bsl-2) laboratory of The Fourth People’s Hospital of Ningxia Hui Autonomous Region.

### Metabolites Extraction

200 μL water was added into EP tube with 10 mg of freeze-dried sample for 30 s vortex. The samples were homogenized at 35 Hz for 4 min and sonicated for 5 min in ice-water bath for 2 times. Then, 480 μL extract solution (MTBE: MeOH= 5: 1) was added into tubes for 30 s vortex, and the samples were further sonicated for 10 min. Then the 300 μL supernatant of total extractions was dried in a vacuum concentrator at 37 °C. Then, the dried samples were reconstituted in 100 μL of 50% methanol in dichloromethane by sonication, and the constitution was then centrifuged at 13000 rpm for 15 min at 4 °C. Fianlly, 75 μL of supernatant was transferred to a fresh glass vial for LC/MS analysis(21).

### LC-MS/MS Analysis

The UHPLC separation was carried out using a ExionLC Infinity series UHPLC System (AB Sciex), equipped with a Kinetex C18 column (2.1 * 100 mm, 1.7 μm, Phenomen). The mobile phase A consisted of 40% water, 60% acetonitrile, and 10 mmol/L ammonium formate. The mobile phase B consisted of 10% acetonitrile and 90% isopropanol, which was added with 50 mL 10 mmol/L ammonium formate. The analysis was carried with elution gradient as follows: 0∼12.0 min, 40%∼100% B; 12.0∼13.5 min, 100% B; 13.5∼13.7 min, 100%∼40% B; 13.7∼18.0 min, 40% B. The column temperature was 45 °C.The injection volume was 2 μL (pos) or 2 μL (neg), respectively.

The TripleTOF 5600 mass spectrometer was used for MS/MS spectra detection on an information dependent basis (IDA). In this mode, the most intensive 12 precursor ions with intensity above 100 were chosen for MS/MS at collision energy (CE) of 45 eV (12 MS/MS events with accumulation time of 50 msec each). ESI source conditions were set as following: Gas 1 as 60 psi, Gas 2 as 60 psi, Curtain Gas as 30 psi, Source Temperature as 600 °C, Declustering potential as 100 V, Ion Spray Voltage Floating (ISVF) as 5000 V or −3800 V in positive or negative modes, respectively (22).

### Data preprocessing and annotation

An self-built program was developed using R for data analysis. The raw data files (.wiff format) were converted to files in mzXML by ‘msconvert’ program from ProteoWizard (version 3.0.19282).Then, the mzxML files were loaded into Lipid Analyzer for data processing. Peak detection was first applied to the MS1 data. The CentWave algorithm in XCMS was used for peak detection, With the MS/MS spectrum, lipid identification was achieved through a spectral match using self-built MS/MS spectral library(23).

### Metabolomic statistical analyses

Packages Metaboanalyst (v3.0) running on the platform **R** (v2.14.0) were used for statistical analyses based on lipids peak areas (as representative of concentration) (16). These data were scaled using auto scaling mode of software. Differences between MB-infected and MTB-infected groups were compared using an independent t test. The PCA and OPLS-DA analysis were also constructed by Metaboanalyst software, while the VIP value of each identified lipids were also acquired.

### Lipid-related categorizes enrichment analysis

Significant differential lipids (P < 0.05) were matched to Lipid-related categorizes enrichment analysis in Metaboanalyst 3.0. And the Lipid-related categorizes at superclass and subclass level were analyzed. Enrichment P-values were computed from a hypergeometric distribution, and used to identify hub terms.

### Lipid network analysis

To identify the most hub lipids in pathogen-host communication network, correlation- topology network were constructed using R software based on spearman parameter. And the network were visualized and further calculated by cytoscape 3.0, and the degree value of each lipids were used to establish lipid importance in the network. The lipids with variable importance in the projection (VIP) scores >1.0, Log1.5 foldchange > 1.0 and P < 0.05 were used to construct the network.

## RESULTS

### Differences in lipid metabolisms between microbial MTB and MB

To investigate the differences in lipid metabolisms of both MTB and MB, both microbial with three biological replicates were analyzed by a in-depth lipid metabolomics. As a results, 12039 signals from negative and positive mode were detected, and 2877 lipids were identified based on the self-built lipid database. To explore an immediate overview of the patterns in the metabolic data and, a Principle Components analysis (PCA) was performed on the metabolic profiles of both microbial and results are shown in Fig. 1A. The results indicated that the top two main components could explain 63.2% of the variations in the data (Figure 1A). All replicates of MTB clearly separated from those replicates of MB, indicating that the lipid metabolism of MTB were significantly different with MB. And the differences may be concentrated in a few highly affected lipids, or may be present in global lipid metabolism. Furthermore, unsupervised hierarchical clustering, an unsupervised method, was performed to identify the variances of lipid metabolisms between both MTB and MB (Figure 1B). The results reached a consensus with the PCA plots that all replicates of MTB were clustered together and observably separated with the replicates of MB (Figure 1A, B), it means that different microbial are having significantly different characteristics in lipid metabolism. Subsequently, the identified lipids that matched the conditions (log1.5 fold changed ≥ 1; P-value ≤ 0.05, and VIP > 1.0) were defined as differential lipids (Figure 1C). In the MB.vs.MTB pairwise comparison, the level of 170 lipids in MB were higher than that of MTB, whereas 56 lipids were increased in MTB (Figure 1C). Therefore, we concluded that there are significant differences in lipid metabolisms between microbial MTB and MB, leading different host responses based on the host pathogen recognition systems.

**Figure 1.**
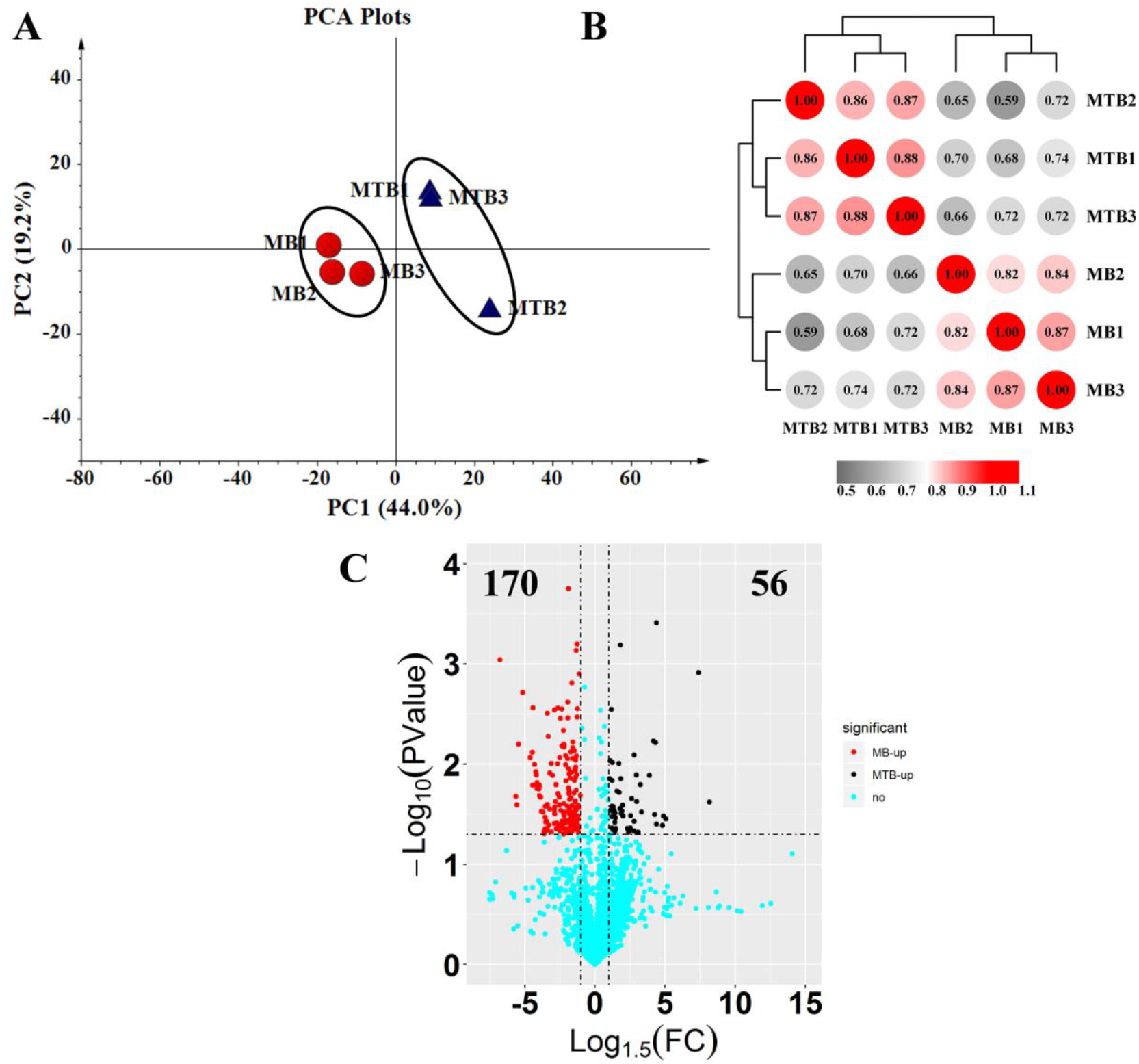
Global analysis of the lipid profile of MTB and MB. (A) Principle component analysis displaying the distinct biological variation among MTB and MB samples. The ellipses with different colors represent the replicates of each MTB or MB sample (red: MB; blue: MTB). All samples are within 95% confidence intervals (Hotelling’s T-squared ellipse). (B) Unsupervised hierarchical clustering of the 6 replicates used in this study showing two distinct clades: MTB and MB. The correlation value between samples were shown in each cell. (C) Volcano plots displayed the differential lipids in MTB vs. MB pairwise comparison. The lipids increased in MB were shown in red, whereas the lipids increased in MTB were shown in black.

### Discovery of hub candidate lipids which caused the differences in lipid metabolism among MTB and MB

To clearly analyze the interference degree among lipid profiles of MTB and MB, the OPLS model was utilized (Figure 2A, B). Each clustering of the OPLS score plot represented a corresponding lipids pattern in MTB or MB. Clearly separation of the MTB.vs.MB pairwise comparison was observed (Figure 2A). The original model R2Y values of the permutation test of the OPLS-DA model was 0.979 (P ≤ 0.05) and very close to 1, demonstrating that the model conforms to the real situation of samples (Figure 2B). Meanwhile, the regression line showed that the Q values in the random model o were all smaller than that of the original model (Figure 2B). Moreover, in the random model, the Q value gradually decreased with the increase in the proportion of Y variable, indicating that the original model is robust and does not over fit (Figure 2B). Overall, all these results above suggested a high model reliability and significant differences in lipid metabolisms among MTB and MB, and the differential lipids identified in this model could well explain the differences in lipids pattern among both MTB and MB.

**Figure 2.**
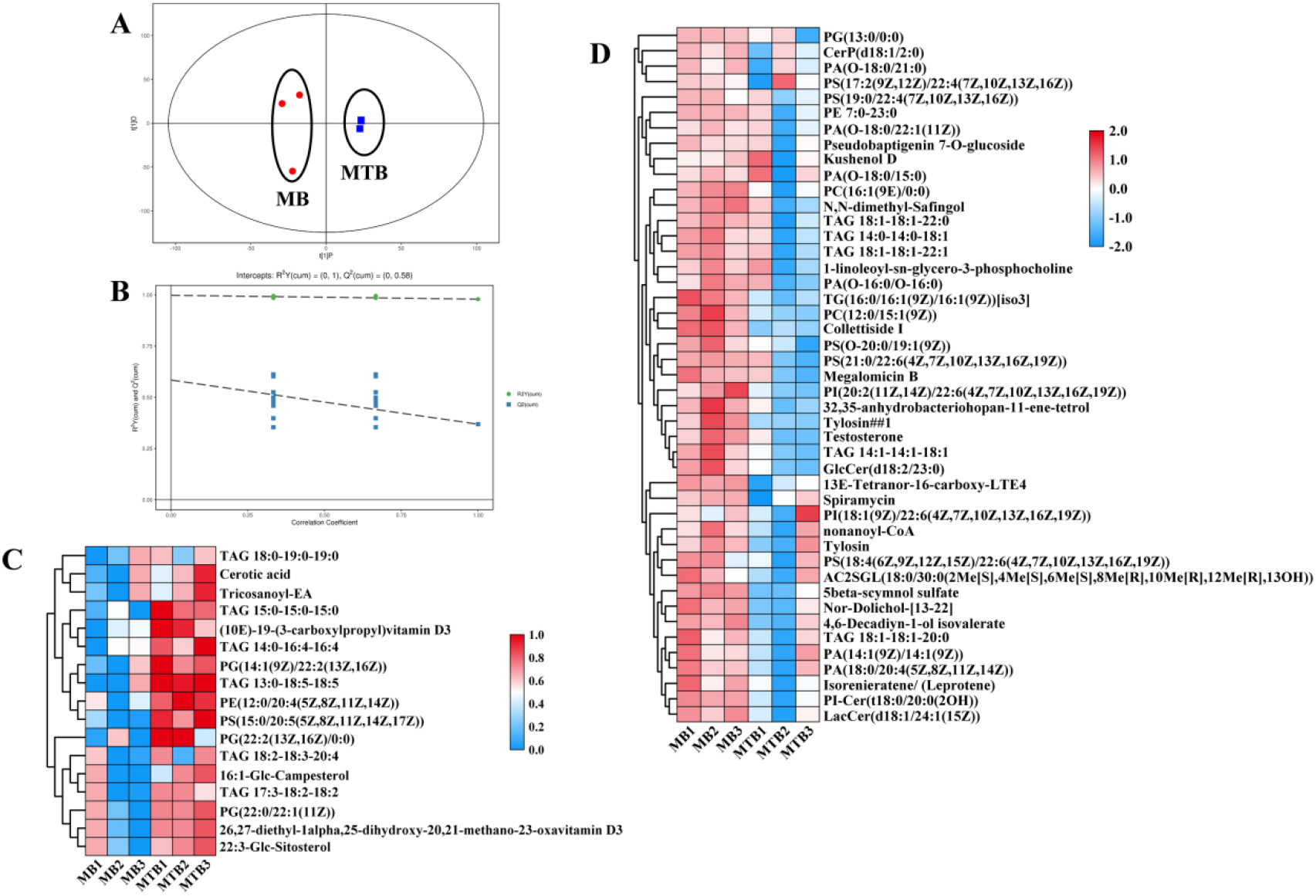
Lipid metabolomics analysis identified specific lipids of MTB and MB. (A)OPLS-DA score plots of MTB and MB showed clear distinction. (B) Permutation test of the OPLS-DA model for MTB vs. MB pairwise comparison. The vertical coordinate represents the value of R2Y or Q with the regression lines of R2Y and Q, and the green dot and blue square shown the value of R2Y and Q obtained by the substitution test, respectively. (C) Heatmap representing the differential lipids that were significantly higher (p < 0.05) in MTB or MB. Data were normalized and auto scaled.

Based on the VIP value of each lipids from OPLS-DA model, 62 lipids were identified as candidate compounds with VIP > 1.0 (Figure 2C, D). And the level of 17 lipids were higher in MTB than that in MB, including PS(15:0/20:5(5Z,8Z,11Z,14Z,17Z)), TAG 13:0-18:5-18:5, PG(14:1(9Z)/22:2(13Z,16Z)), PE(12:0/20:4(5Z,8Z,11Z,14Z)), TAG 15:0-15:0-15:0, (10E)-19-(3-carboxylpropyl)vitamin D3 and TAG 14:0-16:4-16:4 (Table 1; Figure 2C). In parallel, the level of forty-five lipids were higher in MB, such as nonanoyl-CoA, PA(18:0/20:4(5Z,8Z,11Z,14Z)), Nor-Dolichol-[13-22], TG(16:0/16:1(9Z)/16:1(9Z))[iso3], PC(16:1(9E)/0:0), 4,6-Decadiyn-1-ol isovalerate, TAG 14:0-14:0-18:1, TAG 18:1-18:1-22:0, LacCer(d18:1/24:1(15Z)), PC(12:0/15:1(9Z)), GlcCer(d18:2/23:0), TAG 18:1-18:1-22:1 and TAG 14:1-14:1-18:1 (Table 1; Figure 2D). And the top ten candidate lipids of MTB and MB based on their VIP values were shown in Table 1. Five lipids PS(15:0/20:5(5Z,8Z,11Z,14Z,17Z)) , TAG 13:0-18:5-18:5, PG(14:1(9Z)/22:2(13Z,16Z)), PE(12:0/20:4(5Z,8Z,11Z,14Z)) and TAG 15:0-15:0-15:0 were identified as key compounds of MTB lipids metabolism. The level of PS(15:0/20:5(5Z,8Z,11Z,14Z,17Z)) , TAG 13:0-18:5-18:5, PG(14:1(9Z)/22:2(13Z,16Z)), PE(12:0/20:4(5Z,8Z,11Z,14Z)) and TAG 15:0-15:0-15:0 in MTB were 1.61-, 20.02-, 2.18-, 1.70- and 5.62-fold higher than that in MB, with VIP value of 2.18, 2.06, 2.03, 2.02 and 1.88, respectively (Figure 2C; Table 1). And lipids nonanoyl-CoA, PA(18:0/20:4(5Z,8Z,11Z,14Z)), Nor-Dolichol-[13-22], TG(16:0/16:1(9Z)/16:1(9Z))[iso3] and PC(16:1(9E)/0:0) were also identified as key compounds of MB that the level of nonanoyl-CoA, PA(18:0/20:4(5Z,8Z,11Z,14Z)), Nor-Dolichol-[13-22], TG(16:0/16:1(9Z)/16:1(9Z))[iso3] and PC(16:1(9E)/0:0) in MB were 1.57-, 1.75-, 1.90-, 2.08- and 2.02-fold higher than that in MTB, with all VIP value > 2.0 (Figure 2D; Table 1).

**Table 1.** List of candidate differential lipids in MTB or MB

| Lipids name | MTB | MB | VIP | P value |
| --- | --- | --- | --- | --- |
| PS(15:0/20:5(5Z,8Z,11Z,14Z,17Z)) | 0.0000380 | 0.0000236 | 2.175 | 0.003 |
| TAG 13:0-18:5-18:5 | 0.0000043 | 0.0000002 | 2.062 | 0.001 |
| PG(14:1(9Z)/22:2(13Z,16Z)) | 0.0000204 | 0.0000093 | 2.029 | 0.029 |
| PE(12:0/20:4(5Z,8Z,11Z,14Z)) | 0.0000679 | 0.0000399 | 2.017 | 0.027 |
| TAG 15:0-15:0-15:0 | 0.0001190 | 0.0000212 | 1.882 | 0.032 |
| (10E)-19-(3-carboxylpropyl)vitamin D3 / (10E)-19-(3-carboxylpropyl)cholecalciferol | 0.0000456 | 0.0000167 | 1.819 | 0.046 |
| TAG 14:0-16:4-16:4 | 0.0000532 | 0.0000258 | 1.757 | 0.028 |
| PG(22:2(13Z,16Z)/0:0) | 0.0000487 | 0.0000169 | 1.452 | 0.022 |
| 16:1-Glc-Campesterol | 0.0000100 | 0.0000030 | 1.315 | 0.013 |
| 26,27-diethyl-1α,25-dihydroxy-20,21-methano-23-oxavitamin D3 | 0.0000050 | 0.0000018 | 1.270 | 0.033 |
| nonanoyl-CoA | 0.0000047 | 0.0000075 | 2.272 | 0.001 |
| PA(18:0/20:4(5Z,8Z,11Z,14Z)) | 0.0000407 | 0.0000711 | 2.158 | 0.009 |
| Nor-Dolichol-[13-22] | 0.0000078 | 0.0000147 | 2.146 | 0.009 |
| TG(16:0/16:1(9Z)/16:1(9Z))[iso3] | 0.0000081 | 0.0000168 | 2.142 | 0.022 |
| PC(16:1(9E)/0:0) | 0.0000137 | 0.0000277 | 2.141 | 0.008 |
| 4,6-Decadiyn-1-ol isovalerate | 0.0000026 | 0.0000056 | 2.138 | 0.003 |
| TAG 14:0-14:0-18:1 | 0.0000901 | 0.0001540 | 2.136 | 0.008 |
| TAG 18:1-18:1-22:0 | 0.0000075 | 0.0000145 | 2.114 | 0.030 |
| LacCer(d18:1/24:1(15Z)) | 0.0000058 | 0.0000141 | 2.106 | 0.006 |
| PC(12:0/15:1(9Z)) | 0.0000189 | 0.0000333 | 2.099 | 0.044 |

"P < 0.05, VIP > 1.0"

### The lipids composition of MTB and MB were different with each other

The identified candidate lipids in MTB or MB were catagorized by lipid enrichment analysis using MetaboAnalyst 3.0(16). Both level of lipid superclass and subclass were analyzed, respectively (Figure 3). At the superclass level of lipids categorize, enrichment analysis of 17 significant lipids that highly increased in MTB indicated significance in the Glycerophospholipids, Sterol Lipids and Fatty Acyls terms (P < 0.05; Figure 3A). And Glycerophospholipids were the most markedly lipid terms in MTB, with the P value of 0.003 (Figure 3A). Further enrichment analysis of these 17 lipids at subclass level identified their enrichment in terms relevant to Diacylglycerophosphoglycerols, Monoacylglycerophosphoglycerols, Saturated Fatty Acids, N-acyl ethanolamines, Stigmasterols, Ergosterols, Diacylglycerophosphoserines and Diacylglycerophosphoethanolamines (Figure 3C). It has been reported that Ergosterols were important constituents of microbial cell wall, and could be recognized by host cell recognition system by cell wall receptor to activate the host defense responses (24). Among these eight lipid subclasses, Diacylglycerophosphoglycerols were the most marked subclass in MTB that the content of 4 lipids belonged to this term were higher in MTB than that in MB (Figure 3C).

**Figure 3.**
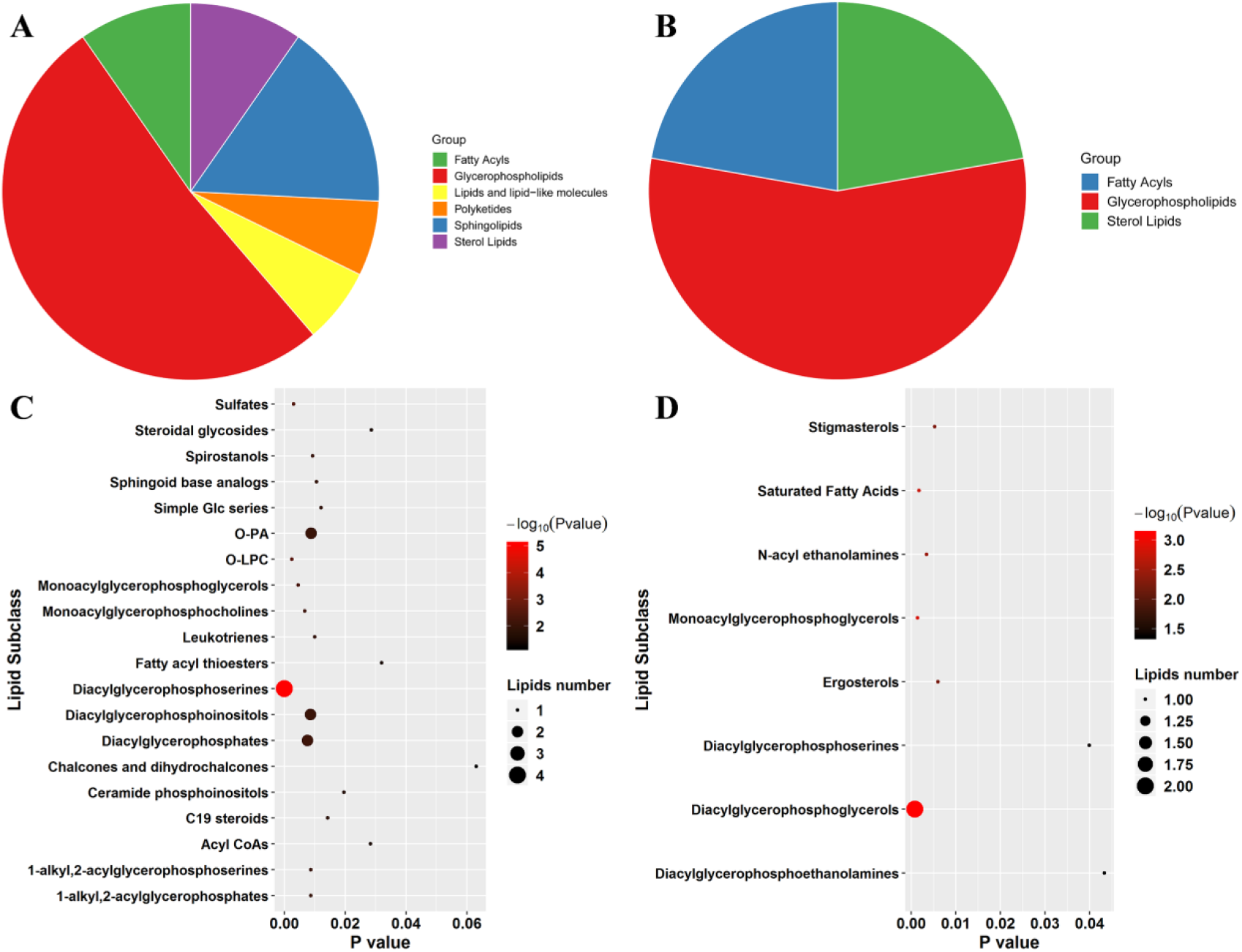
The lipid composition of MTB and MB were significantly different. (A, B) Pie graph displayed the significantly enriched lipid categorizes at superclass level (A: MB group; B: MTB group). (B) Plots depicted the enriched terms relevant to lipid subclass level in MTB and MB. And the plots size represented the number of lipids in the corresponding terms. The vertical coordinate and plots color represent the P-value (−ln(P-value)) of enrichment analysis (C: MB group; D: MTB group).

Subsequently, enrichment Analyses at superclass level of 45 lipids which were higher in MB shown that 6 terms, including Glycerophospholipids, Sterol Lipids, Fatty Acyls, Polyketides, Sphingolipids and Lipids and lipid-like molecules, were significantly enriched among these lipids in MB (Figure 3B). Of note, both MTB and MB were rich in lipids belonged to Glycerophospholipids, Sterol Lipids and Fatty Acyls terms (Figure 3A, B). By contrast, the lipid structure of MB still contain Polyketides, Sphingolipids and Lipids and lipid-like molecules, which were notsignificantly identified in MTB (Figure 3A, B). Furthermore, at the lipid subclass level, we found that the 45 lipids which were rich in MB were mainly belonged to Diacylglycerophosphoserines, O-LPC, Sulfates, Monoacylglycerophosphoglycerols, Monoacylglycerophosphocholines, Diacylglycerophosphates, Diacylglycerophosphoinositols, 1-alkyl,2-acylglycerophosphoserines, 1-alkyl,2-acylglycerophosphates, O-PA and Spirostanols (Figure 3D). Based on this we concluded that although both MTB and MB are rich in Diacylglycerophosphoserines, the lipid composition of MTB and MB were significantly different with each other.

### MTB and MB challenge induce different alterations in lipid metabolism of host macrophage

Lipids of microbial could be recognized by host cell to invoke the defense-related immune responses(19). Hence, we further detected the lipid profiles of BOVINE ALVEOLAR MACROPHAGEfollowing MTB and MB attacks (MTBA and MBA) to investigate whether both microbial with different lipid structure could induce variations in lipid metabolism of BAM. Principal Component Analysis (PCA) on all detected features (Figure 4A) unveiled observably clustering and separation of both MTB- and MB-infected BOVINE ALVEOLAR MACROPHAGEgroups included in this experiment that the BOVINE ALVEOLAR MACROPHAGEfollowing MTB challenge in the negative sector of PCA plots, whereas the BOVINE ALVEOLAR MACROPHAGEattacked by MB was found in the positive sector of the PCA (Figure 4A). With PC1 and PC2 of PCA analysis explaining 76.2% variance among both experiment groups (Figure 4A), the results suggested that MTB and MB challenge induced different alterations in lipid metabolism of BAM. Similarly, correlation analysis on the lipid profiles from these samples also displayed a same results that MTB and MB attacks resulted in different variances in lipid metabolism of BAM(Figure 4B).

**Figure 4.**
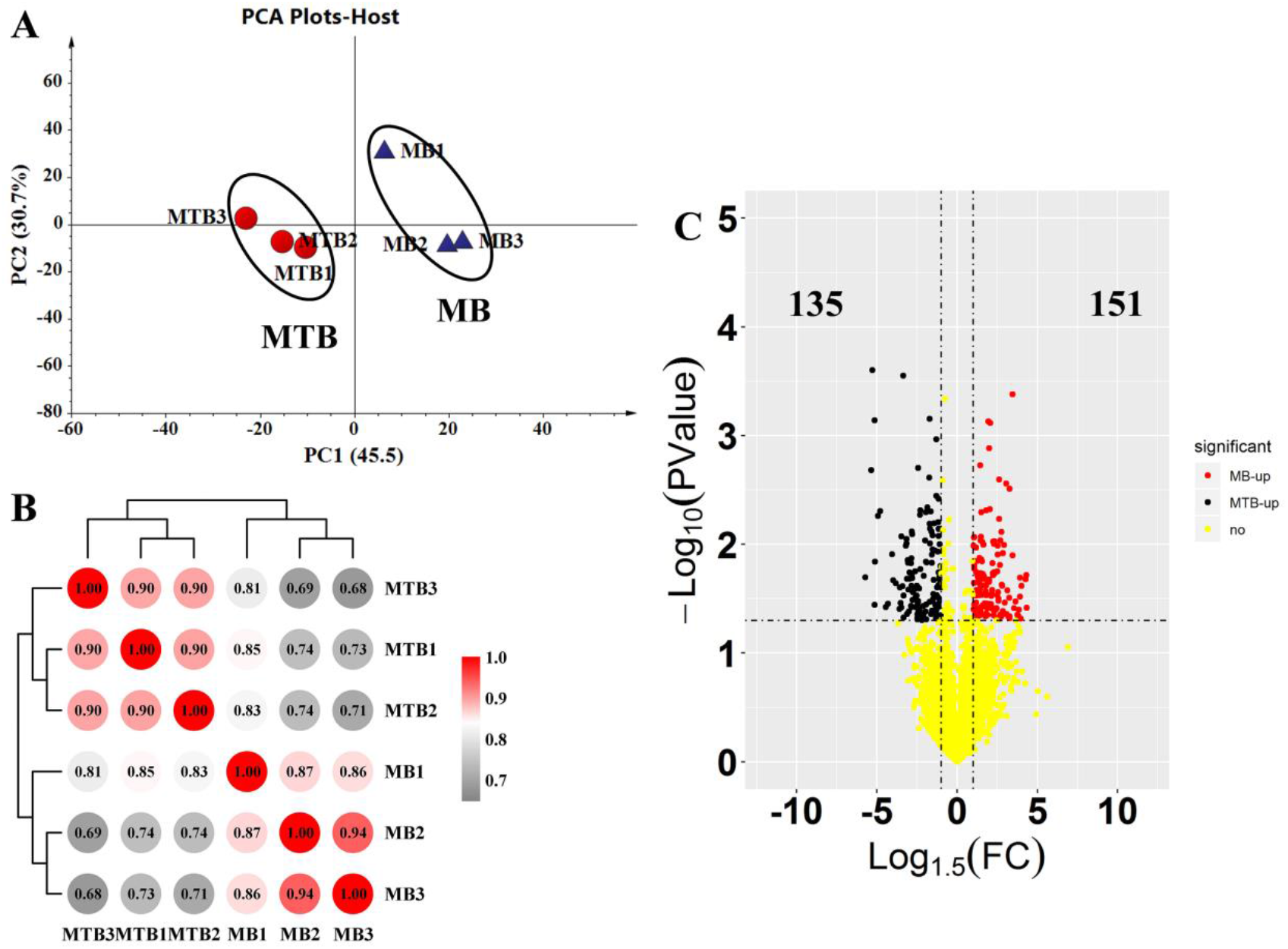
Landscape analysis of the lipid profile of bovine alveolar macrophage following MTB and MB attacks. (A)Principle component analysis on the lipid data of bovine alveolar macrophage attacked by MTB and MB. Two regions within 95% confidence intervals (Hotelling’s T-squared ellipse) were constructed in the PCA plots, including MTBA and MBA which were shown in red and purple, respectively. (B) Unsupervised correlation analysis showing two distinct clades: samples of MTBA and MBA. The correlation value between samples were shown in each cell. (C) Volcano plots showed the differential lipids in MTBA vs. MBA comparison. The lipids increased in bovine alveolar macrophage attacked by MTB were shown in black, whereas the lipids increased in bovine alveolar macrophage attacked by MB were shown in red. The yellow points represented the features with no significance.

Same as the method above, the lipids match the condition (Log1.5 Foldchange > 1.0 and P < 0.05) were identified as differential compounds. Comparison of the BOVINE ALVEOLAR MACROPHAGEfollowing MTB attacks with the ones challenged by MB (MTBA .vs MBA pairwise comparison) revealed 286 significantly differential features (Figure 4C). The level of 135 features were higher in MTBA groups, whereas 151 features were highly increased in MBA groups (Figure 4C).

### MTB and MB attacks induced the accumulation of different kinds of lipids in BAM

Among 286 features, 86 lipids were identified against the self-built lipid database. And thirty-seven of 86 identified lipids increased in BOVINE ALVEOLAR MACROPHAGEfollowing MTB attacks, such as Cer-AP t14:0/22:0, HBMP 14:0-18:1-18:1, PI 2:0-16:4, bacteriohopane-,32,33,34-triol-35-(N-(9-cyclohexyl-nonanoyl))-glucosamine and Cuneatin methyl ether (Figure 5A). Remarkably, the level of Cer-AP t14:0/22:0, bacteriohopane-,32,33,34-triol-35-(N-(9-cyclohexyl-nonanoyl))-glucosamine and Cuneatin methyl ether of macrophages following MTB challenges were 2138.52-, 1061.32- and 684.43-fold higher than that of MB challenged BOVINE ALVEOLAR MACROPHAGE(Figure 5A; Table 2). Meanwhile, 49 identified lipids were highly accumulated in MB-infected BAM, such as PI 15:0-18:0, Ximaosteroid D, Cer-AS d22:2/22:2, GlcADG 22:0-18:1 and Hildgardtol A with VIP values > 2.0 (Figure 5B; Table 2).

**Figure 5.**
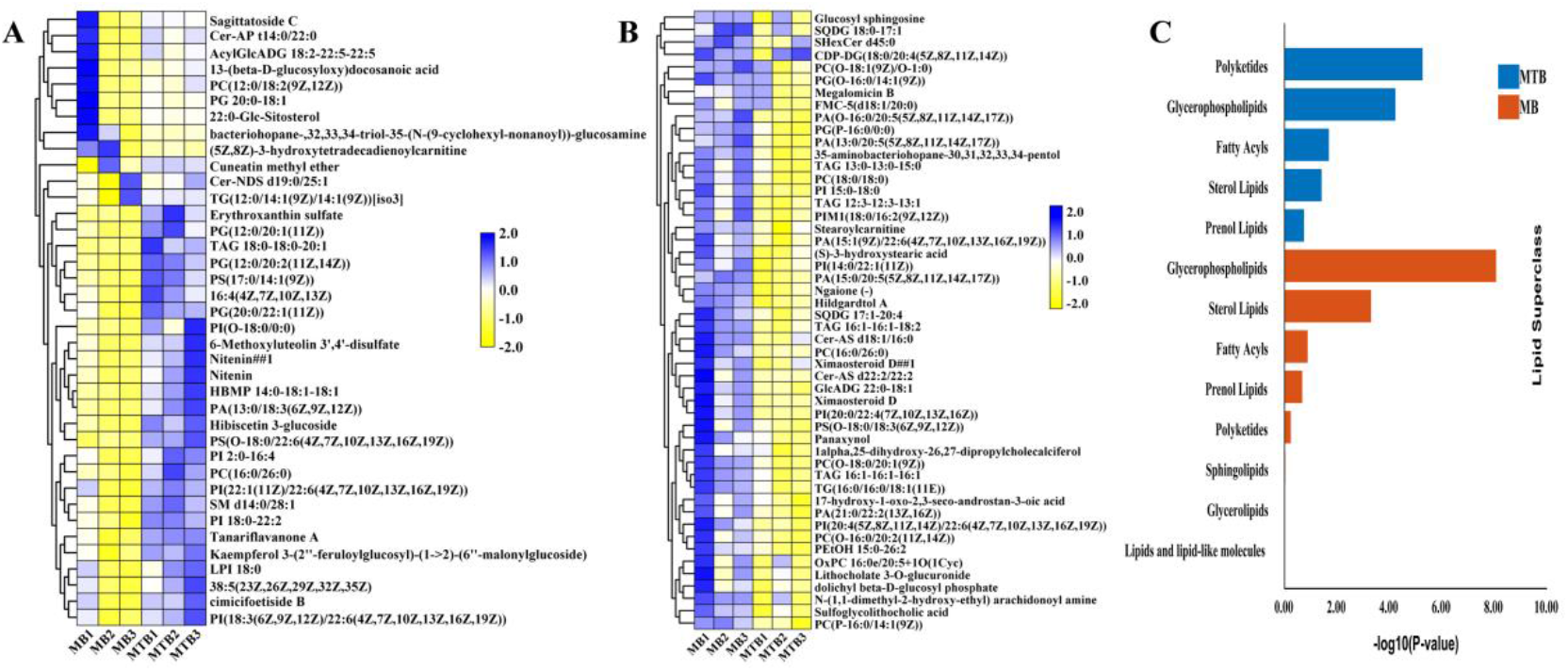
MTB and MB could caused different influences on the lipids metabolisms of BAM. (A, B) Heatmap showed the differential lipids that were significantly higher (p < 0.05) in macrophages attacked by MTB or MB. Data were normalized and auto scaled based on the relative lipid concentration. (C) Bar diagram displayed the significantly enriched lipid categorizes at superclass level. And the terms in MTB- or MB-infected groups were shown in blue and orange, respectively.

**Table 2.** List of candidate differential lipids in macrophages following MTB or MB attacks

| MS2 name | VIP | Pvalue | Foldchange<br>(MBA.vs.MTBA) |
| --- | --- | --- | --- |
| Cer-AP t14:0/22:0 | 2.43 | 0.01 | 0.00047 |
| HBMP 14:0-18:1-18:1 | 2.34 | < 0.01 | 0.4722 |
| PI 2:0-16:4 | 2.33 | 0.01 | 0.55947 |
| bacteriohopane-,32,33,34-triol-35-(N-(9-cyclohexyl-nonanoyl))-glucosamine | 2.33 | < 0.01 | 0.00094 |
| Cuneatin methyl ether | 2.33 | < 0.01 | 0.00146 |
| Nitenin | 2.28 | 0.01 | 0.27954 |
| SM d14:0/28:1 | 2.28 | 0.01 | 0.50515 |
| TAG 18:0-18:0-20:1 | 2.19 | 0.02 | 0.28087 |
| PG(12:0/20:2(11Z,14Z)) | 2.19 | 0.01 | 0.13522 |
| PA(13:0/18:3(6Z,9Z,12Z)) | 2.19 | < 0.01 | 0.14251 |
| PI 15:0-18:0 | 2.3 | 0.02 | 2.47879 |
| Ximaosteroid D | 2.3 | 0.01 | 1.81835 |
| Cer-AS d22:2/22:2 | 2.28 | 0.01 | 1.5474 |
| GlcADG 22:0-18:1 | 2.2 | 0.01 | 2.53818 |
| Hildgardtol A | 2.19 | 0.01 | 3.16664 |
| PI(20:0/22:4(7Z,10Z,13Z,16Z)) | 2.18 | 0.01 | 2.43562 |
| (S)-3-hydroxystearic acid | 2.17 | 0.01 | 3.05692 |
| TAG 12:3-12:3-13:1 | 2.16 | 0.02 | 2.76253 |
| PC(O-16:0/20:2(11Z,14Z)) | 2.14 | 0.01 | 1.90056 |
| PI(20:4(5Z,8Z,11Z,14Z)/22:6(4Z,7Z,10Z,13Z,16Z,19Z)) | 2.13 | 0.01 | 1.61051 |

“P < 0.05 and VIP > 2.0”

Subsequently, the identified candidate lipids in MTBA or MBA were catagorized by lipid enrichment analysis using MetaboAnalyst 3.0(16). At the superclass level of lipids categorize, the enrichment analysis shows the lipids increased in MTBA groups were mainly categorized into terms including Polyketides, Glycerophospholipids, Fatty Acyls, Sterol Lipids and Prenol Lipids (Figure 5C). And the Polyketides and Glycerophospholipids terms were markedly influenced by MTB attacks, with the smallest P value < 0.0001 (Figure 5C). Furthermore, Figure 5C shows the enrichment of Glycerophospholipids, Sterol Lipids, Fatty Acyls, Prenol Lipids, Sphingolipids, Glycerolipids and Lipids and lipid-like molecules terms in the MB challenged group (MBA; Figure 5C). Conversely, MB attacks did not significantly affected the Polyketides-related lipids of BAM. Of note is that MB dramatically induced the accumulation of Glycerophospholipids-related lipids in BOVINE ALVEOLAR MACROPHAGEthat 18 Glycerophospholipids-related lipids were highly increased in BOVINE ALVEOLAR MACROPHAGEfollowing MB infection (Figure 5C). Therefore, the challenges of MTB or MB with different lipids induced different alterations in lipid metabolisms of BAM, supporting our hypothesis above.

### Discovery of hub lipids in the communication network of host macrophages and microbial MTB or MB

To investigate which lipids of microbial caused the alterations in lipids metabolisms of macrophages, all 62 identified lipids of microbial MTB and MB together with 86 lipids identified in macrophages following MTB or MB attacks were used to construct the correlation network using cytoscape 3.0 based on the spearman mode (Figure 6). Using this method, we generated a lipids network of macrophages associated with lipids composition of MTB and MB (Figure 6). This network highlights proposed links between identified lipids from the host macrophages and microbial MTB and MB. In this lipids network which connected lipid structure of the host and pathogens, we used the degree parameter (degree value > 15) to represent the importance of lipids in the network (Supplemental Table 1-2). For MTB, we identified TAG 13:0-18:5-18:5 were the hub lipid which significantly affected the lipid metabolisms of macrophages during the infection process of MTB (Supplemental Table 1; Figure 6). Such compound positively and negatively affected the lipids pattern of macrophages, with correlation value > 0.88 (Supplemental Table 2). Of note, TAG 13:0-18:5-18:5 was specifically increased in MTB, and the level of it in MTB were 20.0-fold higher than that in MB, with VIP value > 2.0 (Table 1; Figure 2). In parallel, four lipids PC(16:1(9E)/0:0), PI(20:2(11Z,14Z)/22:6(4Z,7Z,10Z,13Z,16Z,19Z)), 4,6-Decadiyn-1-ol isovalerate and LacCer(d18:1/24:1(15Z)) of MB were identified as hub compounds in this network, with degree value > 15 (Supplemental Table 1; Figure 6). The level of all these four lipids in MB were about 2.0-fold higher than that in MTB, and significantly affected the lipid metabolisms of macrophages during the infection process of MB, with correlation value > 0.88 (Table 1; Figure 2; Supplemental Table 2). Therefore, we highlight these 5 lipids of MTB or MB in the communication network with host macrophages, leading different variations in lipid metabolisms of macrophages in response to MTB or MB attacks.

**Figure 6.**
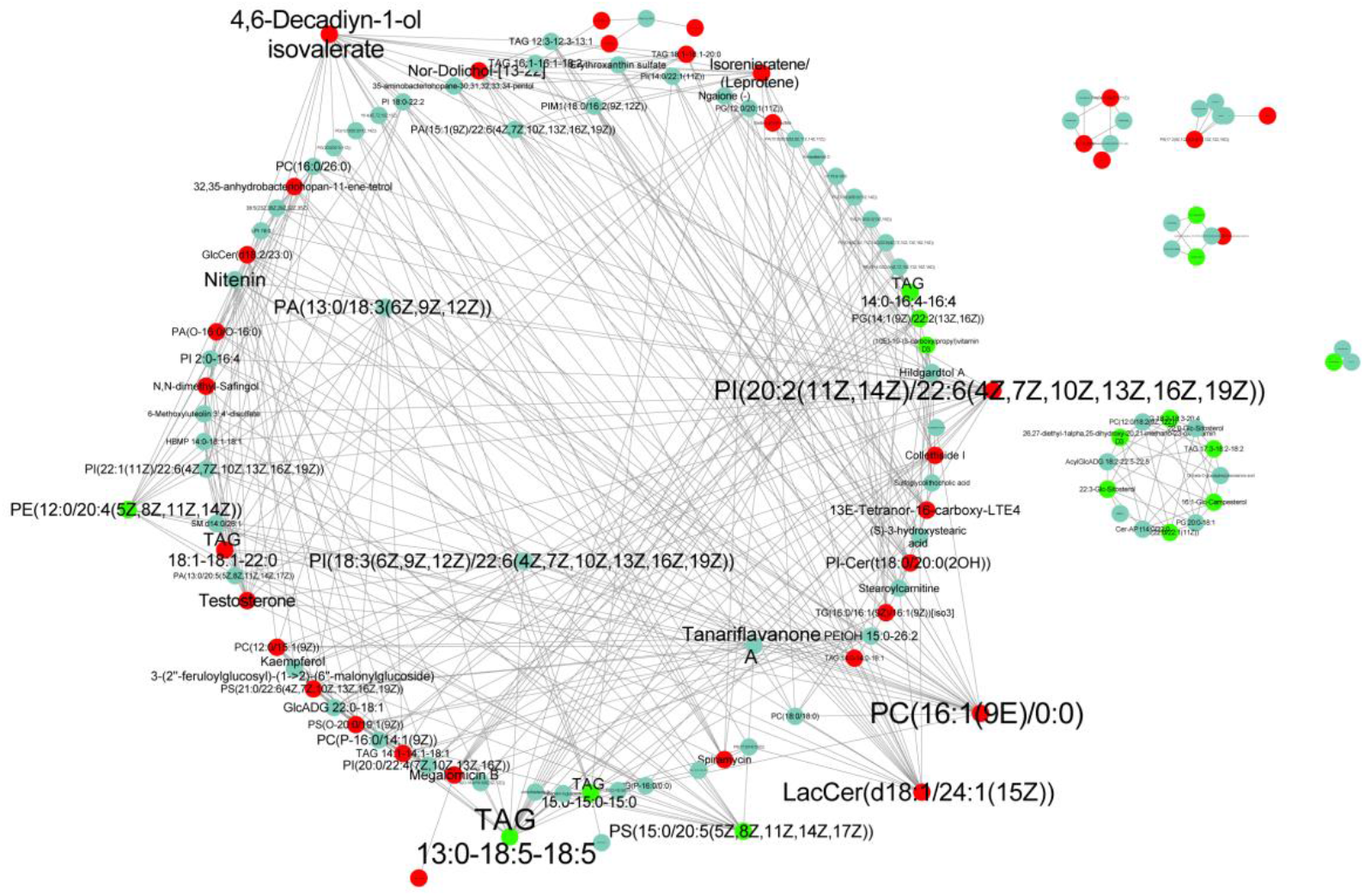
Communication network of lipids-coexpression between microbial and macrophages following microbial attacks. Lipid network constructed through correlation analysis based on spearman parameter. Hub lipid in the network of MTB or MB were shown in green and red, respectively. And the size of each plots represented the degree of each lipid in the network. The plots shown in blue represented the lipids significantly differential in macrophages following MTB or MB attacks.

## DISCUSSION

As a notorious pathogenic bacterial causing tuberculosis (TB) in a broad range of mammalian species, especially humans and cattle, MTB and MB seriously endanger the health of humans and cattle and cause tremendous economic losses annually(1–5). In this study, a dual-lipid metabolomics were performed to elucidate the communication of MTB or MB with macrophages in lipid metabolisms. We found significant variances in lipid composition between MTB and MB that the difference of lipid composition between MTB and MB is mainly in Glycerophospholipids, Sterol Lipids, Fatty Acyls, Polyketides, Sphingolipids and Lipids and lipid-like molecules. The level of various lipids relevant to Polyketides, Sphingolipids and Lipids and lipid-like molecules were higher in MB than that in MTB. Additionally, both pathogenic bacteria induced different alterations in lipid metabolisms of macrophages that MTB mainly induced the accumulation of Polyketides- and Glycerophospholipids-related lipids in macrophages, whereas MB mainly invoked increases in content of Glycerophospholipids and Sterol Lipids in macrophages.

Furthermore, we identified TAG 13:0-18:5-18:5 of MTB and PC(16:1(9E)/0:0), PI(20:2(11Z,14Z)/22:6(4Z,7Z,10Z,13Z,16Z,19Z)), 4,6-Decadiyn-1-ol isovalerate and LacCer(d18:1/24:1(15Z)) of MB were the main factors causing the different lipid-related responses of host macrophage to MTB and MB attacks. Our research systemically revealed the differences in host responses to MTB and MB and the variances between MTB and MB which promotes our understanding of the mechanism of MTBC-host interaction, and thus contributes the control of notorious tuberculosis.

As the main host cell and target cell of MTB/MB, macrophages will actively phagocytosis pathogenic bacteria and make complicated interaction with them in vivo(25–27). During this process, macrophages will inhibit the development of MTB/MB in vivo maingly through autophagy (9). In contrast, as an intracellular parasite, MTB/MB has also evolved a variety of strategies to escape the defense-effect of autophagy, such as down-regulating the expression of autophagy-related factors, inhibiting autophagosome acidification(27, 28). Remarkably, it has been proved that high level of lipid in macrophage will suppress the autophagy responses(29). Our lipid profiles clearly showed an increase in content of lipid in macrophage following MTB/MB attacks, and various lipids relevant to Polyketides, Glycerophospholipids and Sterol Lipids were highly induced to increase in macrophages by MTB and MB, respectively. Various researches indicated that such inhibition effect on macrophage autophagy closely correlated with the level of sterol-related lipids (30). Fatty acid molecules, as ligand of signaling pathway, can affect the immune response of macrophages, and saturated fatty acids such as palmitic acid can inhibit the occurrence of autophagy(31). Of note is that the level of sterol lipid in MB also displayed a high level, indicating that MB may secret sterol lipid in macrophage to increase the sterol lipid content of macrophage to suppress autophagy for their colonization and development. However, we not identified the increase in sterol lipid level in MTB, which indicated a different lipid composition between MTB and MB. While, researchers suggested that in order to survive in macrophages for a long time, MTB also inhibits the process of autophagy through another way that MTB infection can promote the expression of miR144* in macrophages, thus inhibiting the occurrence of autophage(32).

During the invasion process of MTB, MTB can not only cause the occurrence of macrophage autophagy, but also lead to the disorder of intracellular fatty acid metabolism which caused the formation of foam cells (33) . Similarly, our lipid profiles of macrophage following MTB infection showed that various lipids-related metabolism were affected by MTB infection compared with MB infection, leading an increase in level of Polyketides and Glycerophospholipids in macrophage. Studies have shown that MTB infection can induce macrophages to absorb large amounts of lipids, resulting in the formation of foam cells, and the lipid intake is also conducive to the survival and colonization of MTB (15). Kim et al. found that MTB infection stimulates macrophages to absorb more lipids, which were stored in lipid droplets in the form of triacylglycerol and promoted macrophages to form foam cells (34). Induction of the FM formation is one of the main mechanisms which MTB suppresses the defense function of macrophages. By inducing the foaming of macrophages, MTB can not only inhibit the phagocytosis and defense function of macrophages, but also provide a nutrient source for the long-term survival of MTB in vivo(35). MTB can directly ingest the fatty acids stored as lipid droplets in macrophages to maintain its nurtient and energy requirements for its development and successful colonization (8). And the main lipid component in foam cells is free or esterified cholesterol. Normally, cholesterol metabolism in macrophages is a dynamic equilibrium process, which depends on cholesterol intake and outflow. The disorder of lipid load and impaired outflow of cholesterol will cause the cholesterol accumulation and lipid droplets in vivo, which resulted in the formation of foam cells (30, 36). And in order to avoid lipid accumulation, macrophages can regulate intracellular lipid content through lipid autophagy, which involved in the cholesterol metabolism of macrophages and effectively avoided the problem of intracellular lipid accumulation and generation of macrophages (12, 37, 38). However, due to the inhibition of lysosome by MTB, this process cannot be carried out normally. Hence, according to our lipid data we proposed that the infection of MTB is most likely caused the disorder of lipid load and impaired outflow of cholesterol, contributing the formation of foam cells for its development.

In this study, we performed in-depth dual-metabolomics analysis on lipid of the host macrophages and pathogenic MTB and MB. We concluded that MB could through sterol PC(16:1(9E)/0:0), PI(20:2(11Z,14Z)/22:6(4Z,7Z,10Z,13Z,16Z,19Z)), 4,6-Decadiyn-1-ol isovalerate and LacCer(d18:1/24:1(15Z)) to alter the lipid metabolisms of macrophage and suppress its autophagy responses. Meanwhile, MTB could contribute the formation of foam macrophage to successful colonize in host cells. Based on these, we proposed that MTB and MB, with different lipid composition, may suppress macrophage defense responses for their development in host cells through different strategies. However, further in-depth studies on human cells still need to be performed in furture. Overall, We elucidated the lipid communication between host macrophages and MTB and MB with different lipid compositions, which contribute our understanding of the interaction between host macrophage and pathogenic bacteria MTB/MB.

## ACKNOWLEDGMENTS

This study was supported by the Key Project of Research and Development of Ningxia Hui Autonomous Region of China (2017BN04)

